# Empirical evidence of a role for insertion sequences in the repair of DNA breaks in bacterial genomes

**DOI:** 10.1101/2024.06.02.596822

**Authors:** Wing Y. Ngan, Lavisha Parab, Frederic Bertels, Jenna Gallie

## Abstract

**I**nsertion **S**equences (ISs) are mobile pieces of DNA that are widespread in bacterial genomes. IS movements typically involve (i) excision of the IS element, (ii) cutting of the target site DNA, and (iii) IS element insertion. This process generates a new copy of the IS element, as well as a short duplication at the target site. It has been noted that, when observing extant IS element copies in a genome, occasionally no **T**arget **S**ite **D**uplication (TSD) is readily identifiable. This has been attributed to degeneration of the TSD at some point after the insertion event. Here, we provide evidence that some IS movement events – namely, those that occur in association with large-scale genome rearrangements – occur without generating TSDs. In support of this hypothesis, we provide two direct, empirical observations of such IS transposition events: an IS*481* movement occurring with a large duplication in *Pseudomonas fluorescens* SBW25, and an IS*5*/IS*1182* movement plus a large deletion in *Escherichia coli* C. Additionally, we use sequencing data from the Lenski long-term evolution experiment to provide a further 14 examples of IS*150* movements in *E. coli* B that are associated with large deletions and do not carry TSDs. Overall, our results indicate that some IS elements can insert into, and thus repair, existing DNA breaks in bacterial genomes.

## INTRODUCTION

Genomic rearrangements, such as large duplications and deletions, are among the most common types of mutation in bacteria [1–3]. Such rearrangements occur as a result of recombination between distant genomic regions. In bacterial genomes, dispersed copies of mobile genetic elements – such as **I**nsertion **S**equence (IS) and **R**epetitive **E**xtragenic **P**alindromic (REP) sequences – frequently act as hot spots for such recombination events [4, 5]. In the case of ISs, genomic rearrangement is thought to be promoted in at least two ways: (i) by the provision of homologous stretches of DNA that serve as the raw material for recombination, and (ii) through the enzymatic activity encoded within the IS element [6]. The latter mechanism is, as yet, ill-defined.

IS elements, of which there are many families, are the smallest autonomous mobile genetic elements in bacterial genomes [7]. Ranging in size from ∼0.7 kb to ∼2.5 kb, IS elements typically consist of a transposase gene flanked by two, usually inverted, repeats. Most IS elements move and replicate by binding to their own terminal repeats and cutting themselves out of the DNA. The transposase-DNA complex is then inserted elsewhere in the genome, sometimes at specific target sites. The entire transposition process occurs during DNA replication, ensuring that both the original and new IS copies are maintained in the mother and daughter genomes (through DNA repair mechanisms) [8–10]. During insertion, most IS elements generate a **T**arget **S**ite **D**uplication (TSD; sometimes called “direct target repeats”) of between 1-bp and 14-bp, depending on the IS family [10]. These TSDs occur because the transposase cuts the target DNA at different positions on the leading and lagging strands. The resulting overhangs are repaired after the IS element is pasted into the genome, leaving a short, direct repeat at the insertion site.

Through repeated transposition activity, bacterial genomes may carry numerous (nearly) identical, dispersed copies of a particular IS element. Recombination between dispersed IS elements is a common cause for large genomic rearrangements. One example is provided by Lenski’s **L**ong-**T**erm **E**volution **E**xperiment, (LTEE) in which twelve populations of *Escherichia coli* B have been evolving for many thousands of generations [11, 12]. In one evolving population (Ara-1), five of the nine (∼55 %) identified large genomic rearrangements are due to recombination between dispersed IS copies [5]. Similarly, in an experiment with *Salmonella typhimurium*, of 1,800 spontaneous duplication mutants trapped on a plasmid, 97% arose by recombination between flanking IS elements [13]. Intriguingly, when the enzymatic activity of these IS elements was deactivated – either by replacement with two copies of a gene of identical length, or by changing the start codon of the transposase gene – the duplication frequency reduced by two thirds [13]. This result suggests that, in addition to providing the raw material for homologous recombination, transposase activity itself increases recombination rates and hence intra-genomic rearrangement frequencies.

In this work, we present empirical observations that support an active role for IS elements during the formation of genomic rearrangements. We demonstrate and characterize the insertion of new IS element copies in association with (i) a large duplication in *Pseudomonas fluorescens* SBW25, and (ii) a large deletion in *E. coli* C. Furthermore, we present evidence for multiple similar events occurring in the evolving *E. coli* B populations of the LTEE. These rearrangements are not mediated through the recombination of two IS copies. Instead, an IS element is found either separating the two copies of a duplicated DNA sequence, or at the center of a DNA sequence deletion event. Interestingly, these IS element insertions do not leave target site duplications, suggesting that the IS-encoded DNA cleavage function is not involved in this process. Hence, we hypothesize that the IS element is inserted into a DNA strand break that occurs during the resolution of the intragenomic duplication or deletion event.

## RESULTS

### Observation 1: IS activity associated with a large duplication event in *P. fluorescens* SBW25

#### SBW25-Dup-IS carries a new IS element separating two copies of a duplicated DNA sequence

The *P. fluorescens* SBW25 genome contains four dispersed copies of a ∼1.3 kb IS*481*-family element (Figure 1A-B). Each copy consists of (i) a transposase gene flanked on either side by 49-bp inverted repeats, and (ii) an upstream REP sequence. In three copies, the transposase gene and inverted repeats are identical in sequence, while the remaining copy differs at nine positions within the transposase gene. The presence of these multiple highly conserved copies strongly implies (relatively) recent transposition activity. However, to the best of our knowledge, there have been no direct observations of this – or any other – IS element moving within SBW25 populations. Here, we observe an IS*481* transposition event in SBW25.

**Figure 1.**
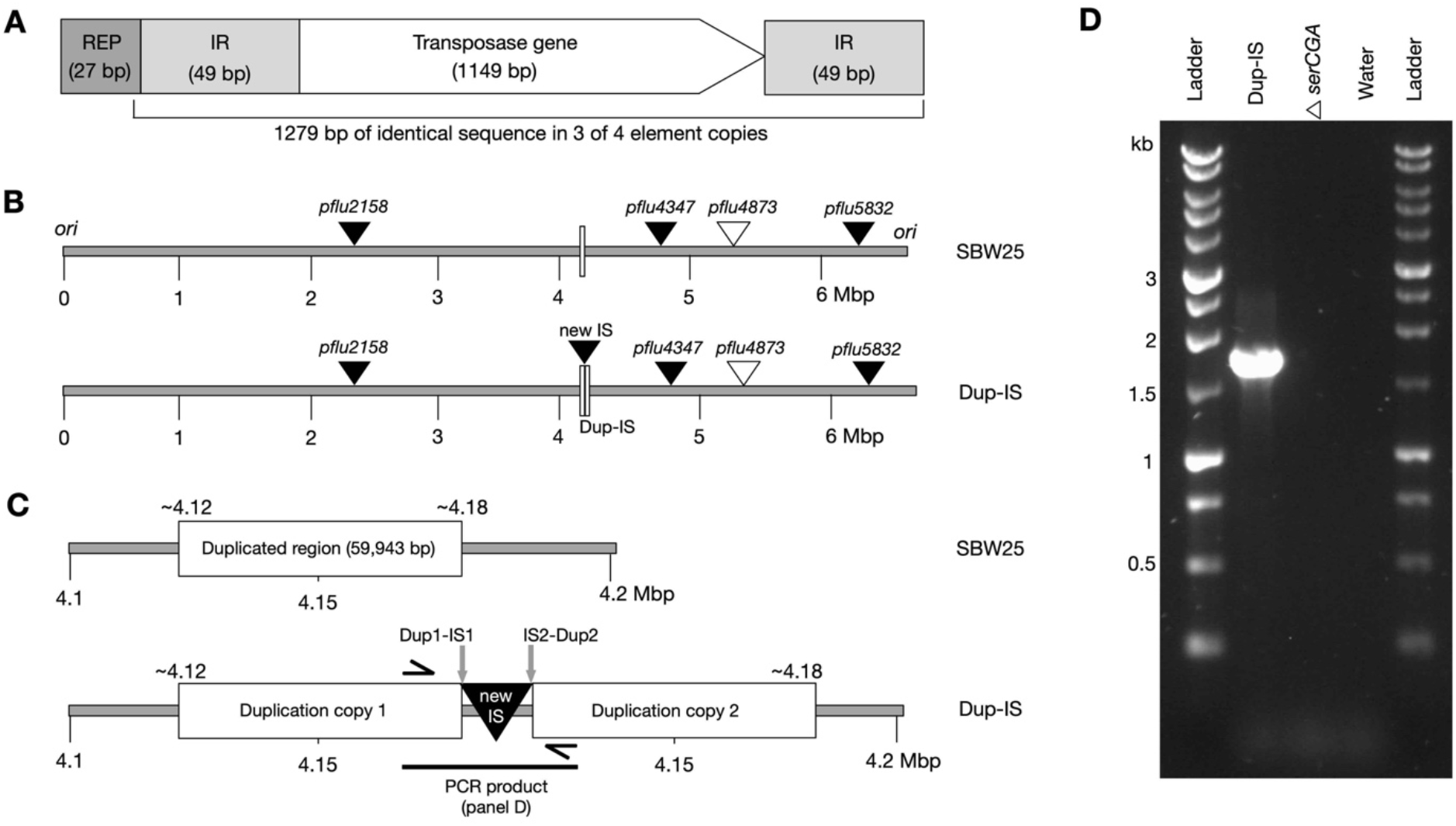
SBW25-Dup-IS carries a genomic rearrangement consisting of a large duplication and an IS*481* transposition event. (**A**) Conserved structure of the SBW25 IS*481-*family transposable element (IR=inverted repeat). (**B**) Linear representations of the SBW25 (top) and SBW25-Dup-IS mutant (bottom) genomes (cut at the origin of replication, *ori*). SBW25 contains three IS*481* copies in which the flanking IRs and transposase genes (*pflu2158, pflu4347, pflu5832*; solid triangles) are identical in sequence, and a fourth copy carrying eight SNPs and a 1-bp deletion in the transposase gene (*pflu4873*; open triangle). The 1-bp deletion is predicted to cause truncation of the transposase protein (R347G*), presumably rendering Pflu4873 enzymatically inactive. SBW25-Dup-IS contains a new, fifth IS copy (solid triangle) and a large duplication (open rectangles). (**C**) Cartoon enlargement of the rearrangement locus. In SBW25-Dup-IS, the two 59,943-bp duplication copies are separated by the new 1,279-bp IS*481* copy (giving a total of 61,222-bp extra DNA). Black, one-sided arrows indicate approximate placement of the primers used for the PCR in panel D. (**D**) PCR-mediated amplification of the rearrangement locus in SBW25-Dup-IS. The 1,746-bp PCR product includes the new IS copy and the two emergent, flanking DNA junctions (Dup1-IS1 and IS2-Dup2).

The strain of interest, SBW25-Dup-IS, was isolated from a laboratory population founded by a slow-growing strain (tRNA gene deletion mutant SBW25Δ*serCGA* [14]). Whole genome re-sequencing revealed that, in line with previous strains isolated under similar conditions [14, 15], SBW25-Dup-IS contains a large, intragenomic duplication: a 59,943-bp duplication of genomic positions 4,119,903 – 4,179,845. However, the SBW25-Dup-IS duplication differs from prior reports in that the two copies of the duplicated region are separated by a new, fifth copy of the IS*481* element (Figure 1B-C; Supplementary Table S2). This proposed Dup1-IS-Dup2 genomic arrangement was confirmed in the laboratory by PCR-mediated amplification and Sanger sequencing of the emergent DNA fragment – encompassing the new IS*481* copy and the novel flanking regions – from SBW25-Dup-IS (Figure 1D; Supplementary Text S1).

Elucidation of the precise Dup-IS-Dup nucleotide sequence revealed two points of interest. Firstly, the movement of the IS element did not generate the 4-bp TSD that is observed for the four other SBW25 IS*481* elements. Indeed, no TSD of any size was observed; notably, the 4-bp sequence that we previously mistook for the TSD was copied along with the rest of the insertion sequence, rather than generated at during the downstream insertion event (see Figure 2). Secondly, insertion occurred into a REP sequence. Analysis of the upstream region of the four previously-existing IS*481* copies reveals a REP sequence at the 5’ end of each, suggesting that REPs – of which there are hundreds of copies in the SBW25 genome [16] – can act as IS*481* target sites (Figure 2). The REP sequence associated with the new IS copy differs from the others in that it lies within a cluster of 12 tandemly-repeated Group 1 and Group 3 REP sequences. There are 11 similar clusters of 4-12 tandemly repeated Group 1 and Group 3 REP sequences dispersed around the SBW25 chromosome [17, 18]. It has previously been hypothesized [17, 18] and empirically observed [14, 15] that these REP clusters can serve as recombination hot spots, resulting in large-scale genomic rearrangements.

**Figure 2.**
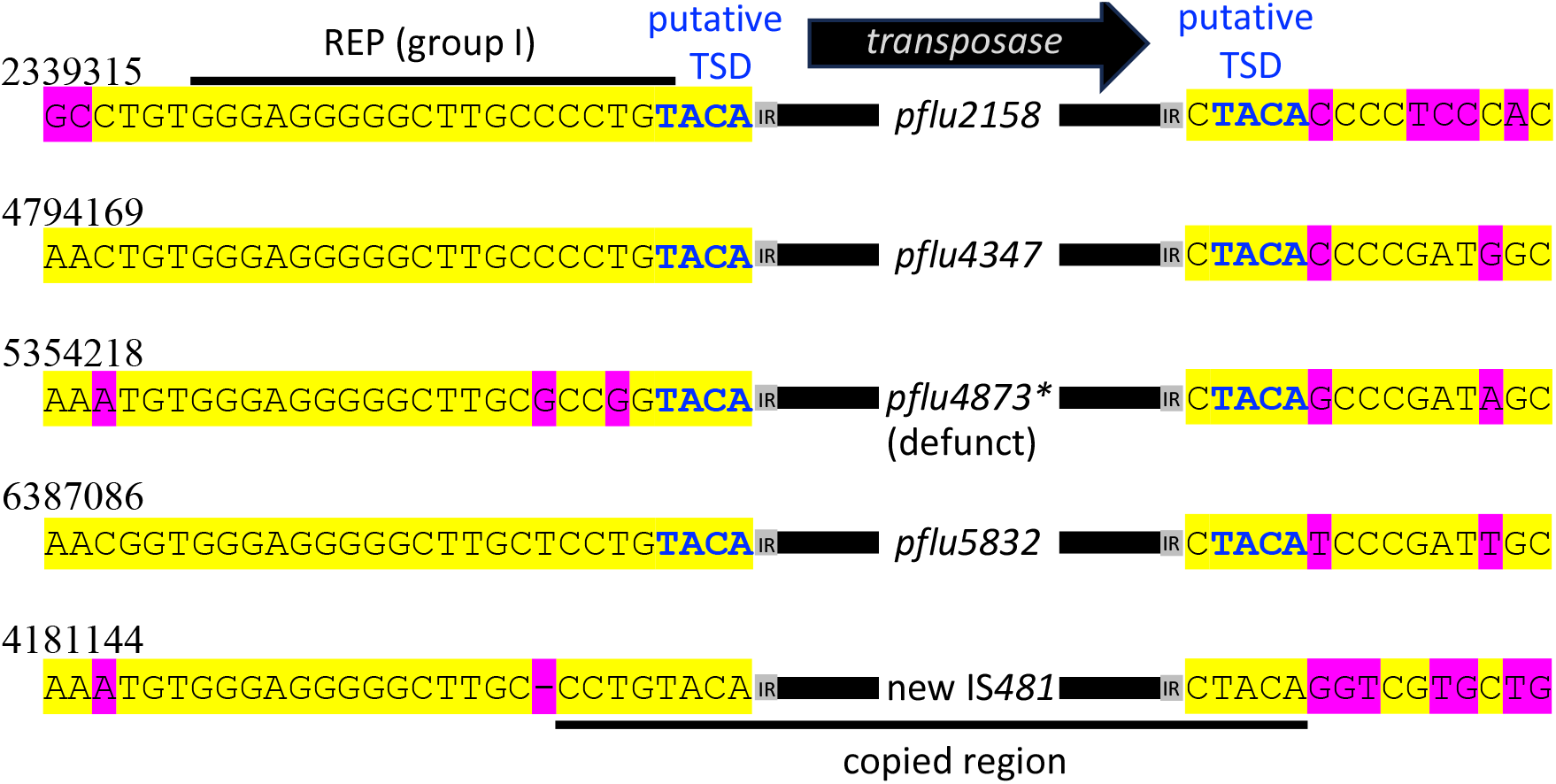
Insertion sites of four established, and one emergent, IS*481* elements in the SBW25-Dup-IS genome. The upstream sequence of each IS*481* insertion site is highly conserved (left; 5’ end of IS). IR=inverted repeat; yellow highlighting=consensus; pink highlighting=deviates from consensus, bold blue text=putative TSDs (note that in the new IS*481* copy no putative TSD exists; see text). In each case, the IS*481* sequence has inserted into the same position of a REP sequence, thereby either (i) disrupting a REPIN (*pflu4347, pflu4873, pflu5832*), (ii) inserting into a REP sequence (*pflu2158*), and/or (iii) inserting into a REP tandem repeat (new IS*481* insertion) [17, 18].

The work in this section characterizes a complex genomic rearrangement event in SBW25-Dup-IS, consisting of (i) the duplication of ∼60 kb genomic DNA, and (ii) the insertion of a new IS*481* copy at the junction of the two duplication fragment copies. The endpoints of the new Dup-IS-Dup genomic arrangement suggest that the duplication event was originally promoted by a tandem REP cluster, but before the duplication could be fully resolved, the IS*481* element inserted into the double-stranded DNA break.

#### The new IS copy, and associated duplication, is readily lost

Above, we demonstrated the emergence of a new, fifth copy of the SBW25 IS*481* element in conjunction with a large duplication. Given that large, tandem duplications are readily lost – usually without a trace – from bacterial genomes [15, 19–21], we wanted to investigate the fate of the new IS copy. That is, if the duplication is lost from SBW25-Dup-IS, does the new IS copy remain?

We began by testing the stability of the SBW25-Dup-IS genome rearrangement with a stability assay [15]. The stability assay involves (i) growing the duplication strain in overnight culture, (ii) plating onto agar, and (iii) examining the size of the resulting colonies. Cells that retain the duplication grow relatively quickly and thus generate large colonies, while cells that have lost the duplication give rise to visibly smaller colonies. On average, 0.91% of colonies derived from overnight SBW25-Dup-IS cultures showed the small colony phenotype, while SBW25Δ*serCGA* (the ancestor of SBW25-Dup-IS) consistently gave rise to colonies of a single size (medians of eight replicates per strain; Figure 3A and Supplementary Table S3). These results indicate that, as expected, the duplication carried by strain SBW25-Dup-IS is unstable.

**Figure 3.**
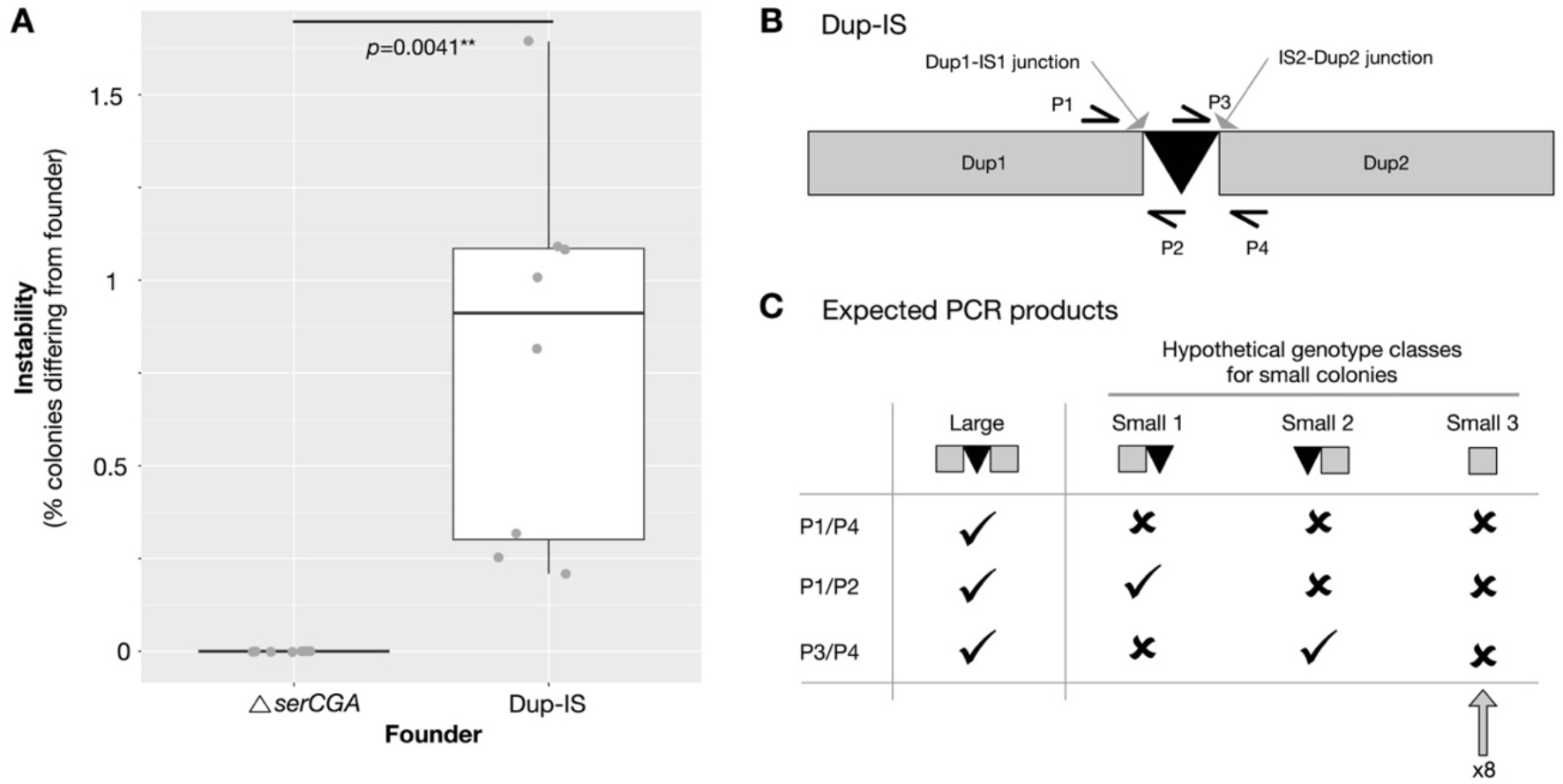
The new IS*481* copy is readily lost from SBW25-Dup-IS, along with the duplication fragment. (**A**) The stability of colony size in ancestral SBW25Δ*serCGA* versus derived strain SBW25-Dup-IS (∼60 kb duplication fragment). Colonies plated from overnight cultures of SBW25Δ*serCGA* consistently retain the SBW25Δ*serCGA* phenotype, while those from SBW25-Dup-IS do not. Eight independent replicates were performed per strain (grey circles), with a minimum of 146 colonies counted per replicate. A Wilcoxon rank sum test was used to test for a difference in median instability (\*\*\**p*<0.001, \*\**p*<0.01, \**p*<0.5). (**B**) Cartoon of SBW25-Dup-IS genotype showing the position of the two emergent junctions (Dup1-IS1 and IS2-Dup2), and the placement of primers to amplify each junction as follows: (i) primers P1/P4 to amplify both junctions and the new IS*481* copy in a single, 1,746-bp product, (ii) primers P1/P2 to amplify Dup1-IS1 (359-bp product), and (iii) primers P3/P4 to amplify IS2-Dup2 (465-bp product). (**C**) Expected PCR products large colonies (i.e., SBW25-Dup-IS) and three hypothetical genotypic classes of SBW25-Dup-IS-derived small colonies: small-1 (retention of IS*481* and the Dup1-IS1 junction), small-2 (retention of IS*481* and the IS2-Dup2 junction), and small-3 (loss of IS*481* and both junctions). All eight small colony isolates show the PCR pattern expected for small-3 (see Supplementary Figure S1 for PCR gels).

SBW25-Dup-IS duplication fragment loss can, hypothetically, result in several new genomic arrangements. These include: (i) loss of the second duplication copy, leaving the new IS element and the first duplication copy (*i*.*e*., emergent junction Dup1-IS1 remains), (ii) loss of the first duplication copy, retaining IS and the second duplication (*i*.*e*., emergent junction IS2-Dup2 remains), and (iii) clean loss of one duplication copy and the IS element (*i*.*e*., neither junction remains). These genomic rearrangements can easily be distinguished from each other using a set of three PCRs. The first PCR, the same as that shown in Figure 1D, is used to demonstrate duplication loss; a product is only amplified in the presence of the SBW25-Dup-IS duplication. In cases where no product is obtained for PCR1, the second and third PCRs indicate whether the new IS element is retained (PCR2 amplifies junction Dup1-IS1, PCR3 junction IS2-Dup2). If the new IS copy remains, a product should be obtained for either PCR2 and PCR3. Alternatively, if the IS element is lost along with the duplication, none of the three PCRs should amplify a product (Figure 3B-C).

To test which type(s) of the above genomic rearrangements are occurring in small colonies arising from SBW25-Dup-IS, sixteen colonies were isolated from the stability assay (one small and one large colony from each of the eight independent replicates). The three PCRs demonstrate that all eight small colonies had lost both the duplication fragment and the new IS element (while all eight large colonies retain both) (Figure 3C; see Supplementary Figure S1 for gels).

### Observation 2: IS activity associated with a large deletion event in *E. coli* C

During a second, independent evolution experiment with *E. coli* C and phage ΦX174, movement of an IS*5*/IS*1182* element was observed in association with a large deletion [22]. Here, we provide an in-depth computational and empirical analysis of this genomic rearrangement. The strain of interest, Ec-Del-IS, contains a 62,007-bp deletion (genomic positions 3,612,369 – 3,674,375) and an insertion of a new IS*5*/IS*1182* element copy at the deletion site (see Supplementary Text S2; Supplementary Table S4). The *E. coli* C IS*5*/IS*1182* element is ∼1 kb in length, almost the entirety of which is a 918-bp transpose gene flanked by 14-bp inverted repeats, without any obvious TSD. Two identical, full copies of the element are present in the wild type *E. coli* C genome (transposase gene locus tags *B6N50_00555* and *B6N50_14295*), and three are found in the Ec-Del-IS genome (Figure 4A-B).

**Figure 4.**
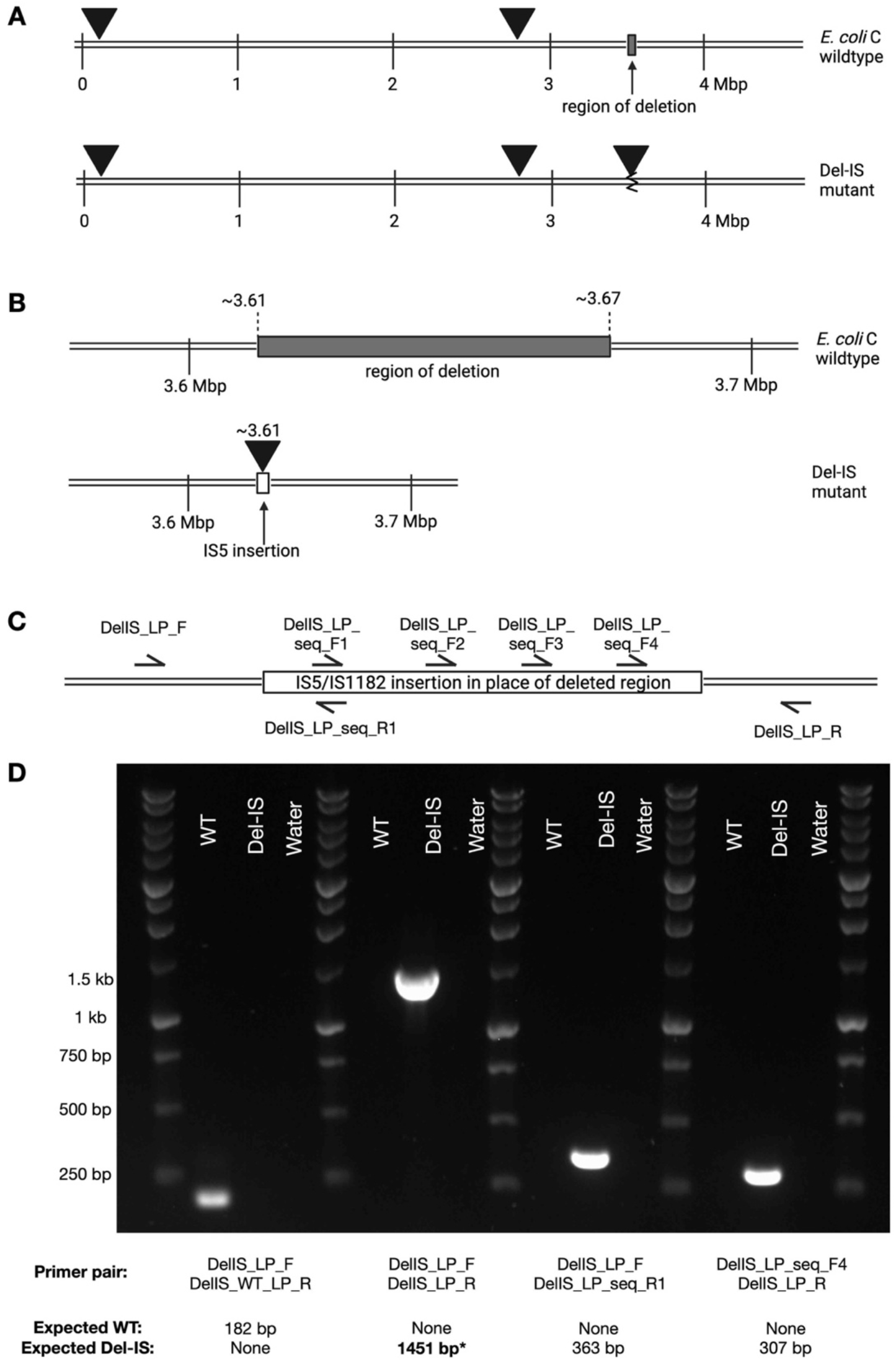
Ec-Del-IS carries a genomic rearrangement consisting of a large deletion and an IS*5*/IS*1182* transposition event. (**A**) A whole-genome view. Linear representations of *E. coli* C wild type (top) and the derived Ec-Del-IS (bottom) (cut at the origin of replication, *ori*). The wild type genome contains two identical copies of the 1,051-bp IS*5*/IS*1182* sequence (*B6N50_00555, B6N50_14295*; solid triangles). The IS mutant contains a ∼62 kb deletion (open rectangle; top) and an extra copy of the IS element (third solid triangle; bottom). (**B**) A close-up view showing the 62,007-bp deletion. (**C**) Cartoon showing positions of the PCR and sequencing primers used to verify the mutation. (**D**) PCR-mediated amplifications of the rearrangement locus in Ec-Del-IS (versus wildtype). In order to confirm the sequence across the genomic rearrangement, the starred 1,451-bp PCR product (which includes the new IS copy and the emergent DNA junctions Del1-IS1 and IS2-Del2) was Sanger sequenced with the seven primers shown in panel C.

The emergence of the deletion-IS mutation in Ec-Del-IS was demonstrated using PCR (Figure 4C-D), and the sequence of the mutated region was verified by Sanger sequencing. No TSDs are observable at the insertion site, suggesting that the IS element insertion occurred at the same time as the deletion event (*i*.*e*., with the IS*5*/IS*1182* element inserting into the double-stranded DNA break). Notably, given the irreversible nature of large deletions, the Ec-Del-IS genotype is far more stable than SBW25-Dup-IS.

### Observation 3: Evidence for frequent IS-Del mutations occurring in the LTEE

So far, we have described two IS movements that (i) are associated with genomic rearrangements (a large duplication in SBW25-Dup-IS, and a large deletion in Ec-Del-IS), and (ii) did not generate the TSDs usually connected with IS movements. In order to assess whether similar IS-rearrangement mutations occur more widely, we turned to publicly available genome sequencing data from the *E. coli* B LTEE experiment. Illumina sequencing reads for three 60,000-generation populations with particularly high IS activity (Ara+1, Ara-3, Ara-6) were analysed with *breseq* [23]. A total of 83 IS movement events were predicted, encompassing four IS families (in decreasing order of prevalence: IS*150* (68), IS*186* (7), IS*1* (5), IS*3* (3)) (Figure 5A; Supplementary Table S5). Of these, 14 (∼17%) occurred in association with a large deletion (of between 14-bp and 48,894-bp), in each case resulting in the same Del-IS-Del configuration as observed in Ec-Del-IS. All 14 simultaneous IS and deletion events involve IS*150* family elements.

**Figure 5.**
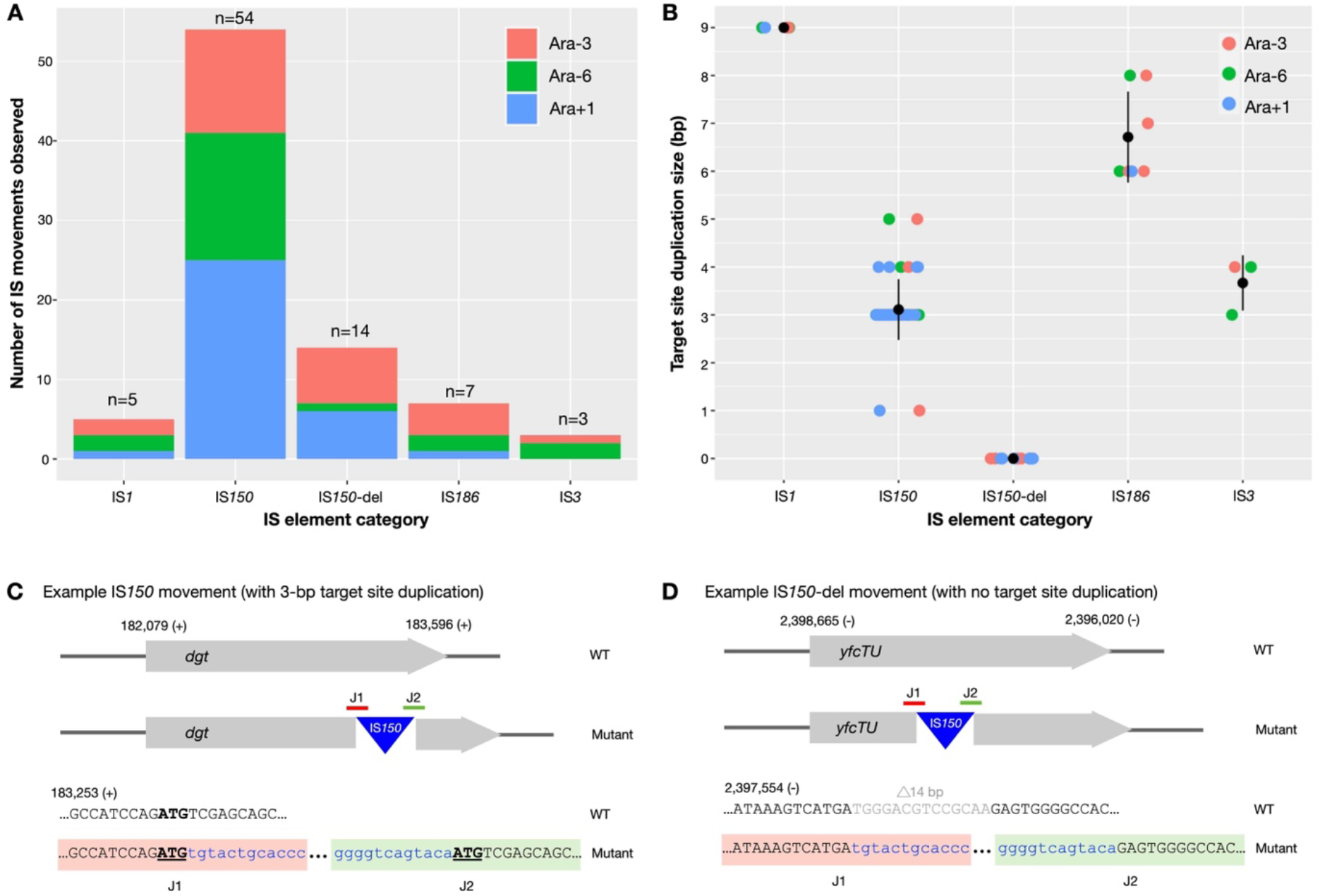
Characterisation of IS transposition events in three LTEE populations at 60,000 generations (Ara+1, Ara-3, and Ara-6). (**A**) 83 distinct IS transposition events were identified, involving IS elements from four families (IS*1*, IS*150*, IS*186* and IS*3*). In the case of the IS*150-*family, the 68 transposition events occurred in two categories: those that are not associated with large deletions (54, category “IS*150*”), and those that are associated with large deletions (14 events, category “IS*150*-del”). (**B**) Across all four IS-families, each of the 69 transposition events not associated with a large deletion generated a TSD (1-bp to 9-bp), while none of the 14 IS*150-*del transposition events generated TSDs. Coloured circles indicate the length of the TSD per transposition event, black circles+lines show the mean+standard deviation for each IS-family. (**C**) An example of an IS*150* insertion event that is not associated with a deletion. The event was identified in Ara+1, and results in the insertion of a new IS*150* element in the *dgt* gene plus the generation of a 3-bp direct repeat at the target site (bold and underlined). Black text=*dgt*, blue text=IS*150*. (**D**) An example of an IS*150-*del insertion event that is associated with a 14-bp deletion. The event was identified in Ara+1, and results in the insertion of a new IS*150* element in the *yfcTU* gene, and the deletion of a 14-bp sequence from *yfcTU* (light grey text). No direct repeats are generated at the target site. Black/grey text=*yfcTU*, blue text=IS*150*.

Having established that two types of IS movements (*i*.*e*., those that occur alone, and those that occur with a genomic rearrangement) exist, we next looked at whether these two transposition categories are likely to occur via distinct molecular mechanisms. To begin, we investigated the identifiable TSDs in each of the IS movement events. As outlined in the introduction, most IS transposition events generate a scar at the target site (short duplications of ∼1-bp to 14-bp). Such scars are due to the transposon enzymatically cutting the target DNA in different positions on the leading and lagging strand during the transposition event. In line with this, all 69 standard IS movements – those that do not coincide with a deletion – observed here generated a TSD of between 1-bp and 9-bp (Figure 5B-C; Supplementary Table S5). Contrastingly, none of the 14 IS movements associated with a large deletion generated a TSD of any size (Figure 5D). This lack of TSDs implies that the deletion-associated IS movements occurred via a novel molecular mechanism, one that does not involve the transposase enzymatically cutting the target DNA.

## DISCUSSION

In this work, we have observed multiple examples of IS elements inserting into large-scale genomic rearrangements. Firstly, we have directly observed two such events occurring in bacterial populations evolving in real-time in the laboratory: an IS*481* movement associated with a large duplication in *P. fluorescens* SBW25, and an IS*5/*IS*1182* movement associated with a large deletion in *E. coli* C. Secondly, we computationally identify a further 14 IS*150* movement events that are associated with large deletions in some LTEE *E. coli* B populations. In addition to occurring in association with a large-scale rearrangement, none of the 16 IS movements presented here generated a TSD. While the transposition process typically generates direct repeats at the insertion site (see Introduction), the existence of (occasional) IS elements with no obvious TSDs has been documented for many years [7]. To date, these missing TSDs have been attributed to loss of the TSD at some point after the IS movement (*e*.*g*., through recombination between two disparate IS copies, or indels/SNPs) [7]. While this may be true in some cases, here we show that *de novo* insertions of IS elements – which have not had time to lose TSDs by subsequent mutation – can also lack TSDs, when they occur in association with large duplications or deletions. We hypothesize that, in these cases, the IS element directly inserts into the double-stranded DNA breaks that occur during the resolution of genomic rearrangements [19]. If the IS element does not cut the DNA itself, then the TSDs that result from staggered cuts on the leading and lagging DNA strands are not expected to be introduced.

We are not the first to propose that transposases can take advantage of DNA strand breaks. In the bacterium *Deinococcus radiodurans*, the HUH transposase IS*DRa2* has been observed to transpose during the reassembly of the genome after gamma irradiation [24]. Gamma rays destroy the genome but also induce the activity of the HUH transposase, which very quickly inserts into the new DNA breaks. Similarly, Tn*7* (encoding transposase TnB) has also been suggested to take advantage of DNA breaks for transposition in *E. coli* [25]. Together with these earlier studies, our work suggests that insertion into DNA breaks may be a broad phenomenon that occurs not only with HUH transposases (such as that encoded within IS*DRa2*), but also with DDE transposases (such as those encoded within IS*481*, IS*5*/IS*1182*, IS*150*, and Tn*7*).

Another example of large rearrangements occurring in conjunction with the insertion of DNA is found in *Helicobacter* and *Neisseria*. In these bacteria, genes encoding restriction modification systems are inserted into the centre of large-scale rearrangements [26]. Bioinformatic analyses of closely related strains have shown that large DNA insertions are often accompanied with large intragenomic duplications or deletions (similar to those observed for the IS movements in our work). The authors hypothesize that these insertions are mediated by the restriction modification enzymes attacking target DNA, and subsequent insertion. Alternatively, one could hypothesize that the DNA has been inserted into existing breaks caused by DNA replication errors.

In eukaryotes, double-stranded DNA breaks can be repaired through a non-homologous end-joining mechanism [27]. This repair mechanism is often accompanied by the insertion of non-specific DNA into the break point [28]. In many cases, retrotransposons are inserted into these DNA breaks [29, 30]. While this opportunistic function may provide short-term benefits, it is presumably detrimental to the host in the long-term, as it significantly increases the likelihood of future transposition events [31]. However, in at least one case, it has been shown that a bacterial Group II intron has been domesticated to function as a DNA repair protein in *Pseudomonas aeruginosa* [32].

Together, our data and analyses suggest that some IS families can use existing DNA breaks – such as those that are generated during the process of resolving deletions or duplications – to insert into new positions in the bacterial genome. This hypothesis is supported by the lack of observable TSDs at the insertion site, suggesting that the DNA break is not generated by the endonuclease function of the transposase itself. Our observations also provide an explanation for the absence of TSDs flanking previously observed IS elements inside bacterial genomes.

## MATERIALS AND METHODS

### Growth conditions

Strains used in this study are listed in Supplementary Table S1. *P. fluorescens* SBW25-derived strains were grown in liquid **K**ing’s Medium **B** (KB; [33]) at 28°C with shaking (∼16 hours). *E. coli* C strains were grown in **L**ysogeny **B**roth (LB) at 37°C with shaking (∼16 hours).

### Genome sequence of SBW25-Dup-IS

SBW25-Dup-IS was sequenced using Illumina 150-bp paired-end reads at the Max Planck Institute for Evolutionary Biology sequencing facility, Germany. Raw reads (deposited on Zenodo) were aligned to the SBW25 wild type genome sequence (NCBI RefSeq NC_012660.1, [16]) using *breseq* [23], giving a mean coverage of 57.6-fold per genomic base. Mutations were identified using a mixture of *breseq*, Geneious (version 2023.2.1), and previously published protocols [14, 15] (see Supplementary Text S1, Supplementary Tables S1 and S2).

### Genome sequence of Ec-Del-IS

This work provides a detailed bioinformatic and empirical analysis of the previously-reported genome sequence of Ec-Del-IS (strain E_coliCRF_phi_GM1_12c1 in [22]). Using *breseq*, raw reads were aligned to an updated version of the standard *E. coli* C genome (GenBank CP020543.1). The updated genome differs by the presence of nine insertions and two substitutions [34]. A mean coverage of 142-fold per genomic base was obtained. Mutations were identified using a mixture of *breseq* [23], Geneious (version 2020.1.2), and previously published protocols [14, 15] (see Supplementary Text S2, Supplementary Tables S1 and S4). Raw sequencing reads, and the updated reference genome, are available on Zenodo.

### PCR and Sanger sequencing

Primers used in this study are listed in Supplementary Table S1. PCRs were performed under standard conditions for High Performance GoTaq® G2 Flexi DNA Polymerase (Promega; M7801), with the addition of 5*x* combined enhancer solution [35] per reaction. PCR products were visualized on 1% agarose gels (90 V, 30-45 minutes) against a 1 kb DNA ladder (Promega). Sanger sequencing was performed by the sequencing unit at the Max Planck Institute for Evolutionary Biology. Raw gel images and Sanger sequencing traces are provided on Zenodo.

### Stability assay

A stability assay [15] was used to test for duplication fragment loss from SBW25-Dup-IS in overnight culture. The SBW25-Dup-IS duplication confers a growth advantage; cells carrying the fragment give rise to large colonies, while those that have lost the duplication generate smaller colonies. Eight replicate colonies of SBW25-Dup-IS and SBW25Δ*serCGA* (a control strain from which only small colonies are expected) were grown to stationary phase in overnight culture (28°C, shaking). Each of the 16 cultures was dilution plated on 1.5% KB agar, and plates were incubated at room temperature (∼20°C) for 48 hours. The numbers and sizes of the resulting colonies were recorded (minimum 146 colonies per replicate; Supplementary Table S3). Per culture, an instability measure - the proportion of colonies differing in size from the original genotype (SBW25-Dup-IS or SBW25Δ*serCGA*) - was calculated. A Wilcoxon rank sum test was used to test for a difference in the median instability measure of SBW25-Dup-IS and SBW25Δ*serCGA* (Figure 3A).

### Movement of IS elements in LTEE populations

Raw sequencing reads for three 60,000-generation LTEE populations (Ara+1, Ara-3, Ara-6) were downloaded from NCBI BioProject database (accession number PRJNA380528) [12]. These three populations were chosen because they show high IS activity [31, 31]. Each set of raw reads was aligned to the *E. coli* B REL606 reference sequence (NCBI RefSeq NC_012967.1) using *breseq* [23] on the polymorphism setting (-p). All predicted IS movement events were recorded (Supplementary Table S5). Next, each predicted IS movement was manually inspected and characterized with respect to (i) association with large genomic rearrangements (defined here as insertions or deletions of >10-bp), and (ii) target site duplications (Supplementary Table S5). IS movements were then classified into five categories: four categories in which the movements were not associated with other large rearrangements (namely, IS*1*, IS*3*, IS*186*, IS*150*) and one category in which the movements were associated with large deletions (IS*150-*del).

## ACKNOWLEDGEMENTS

The authors wish to thank Gunda Dechow-Seligmann for technical assistance.

## AUTHOR CONTRIBUTIONS

Multi-author contributions are listed alphabetically. Conceived and designed the experiments: all authors. Performed the experiments: LP and WYN. Analyzed the data: all authors. Wrote the paper: FB and JG. Commented on the paper: all authors.

## FINANCIAL STATEMENT

This work was supported by the German Research Foundation (DFG) [GA 2895/2-1 to JG], the Max Planck Society [all authors], and the International Max Planck Research School for Evolutionary Biology [LP and WYN].

## DATA AVAILABILITY

All raw data, including NGS data, Sanger sequencing, traces, and gel images are available on Zenodo (https://zenodo.org/records/11402079).

## REFERENCE LIST

1. Anderson RP, Roth JR. Tandem genetic duplications in phage and bacteria. Annu Rev Microbiol 1977;31:473–505.

2. Elliott KT, Cuff LE, Neidle EL. Copy number change: Evolving views on gene amplification. Future Microbiol 2013;8:887–899.

3. Reams AB, Roth JR. Mechanisms of gene duplication and amplification. Cold Spring Harb Perspect Biol 2015;7:a016592.

4. Shyamala V, Schneider E, Ames GF. Tandem chromosomal duplications: role of REP sequences in the recombination event at the join-point. 1990;9:939–946.

5. Raeside C, Gaffé J, Deatherage DE, Tenaillon O, Briska AM, et al. Large chromosomal rearrangements during a long-term evolution experiment with Escherichia coli. mBio 2014;5:e01377–14.

6. Hedges DJ, Deininger PL. Inviting instability: Transposable elements, double-strand breaks, and the maintenance of genome integrity. Mutat Res 2007;616:46–59.

7. Mahillon J, Chandler M. Insertion sequences. Microbiol Mol Biol Rev 1998;62:725–774.

8. Ton-Hoang B, Pasternak C, Siguier P, Guynet C, Hickman AB, et al. Single-stranded DNA transposition is coupled to host replication. Cell 2010;142:398–408.

9. Hagemann AT, Craig NL. Tn7 transposition creates a hotspot for homologous recombination at the transposon donor site. Genetics 1993;133:9–16.

10. Hickman AB, Dyda F. Mechanisms of DNA transposition. Microbiol Spectr 2015;3:3.2.12.

11. Lenski RE, Rose MR, Simpson SC, Tadler SC. Long-term experimental evolution in Escherichia coli. I. Adaptation and divergence during 2000 generations. Evolution 1991;138:1315–1341.

12. Good BH, McDonald MJ, Barrick JE, Lenski RE, Desai MM. The dynamics of molecular evolution over 60,000 generations. Nature 2017;551:45–50.

13. Reams AB, Kofoid E, Kugelberg E, Roth JR. Multiple pathways of duplication formation with and without recombination (RecA) in Salmonella enterica. Genetics 2012;192:397–415.

14. Ayan GB, Park HJ, Gallie J. The birth of a bacterial tRNA gene by large-scale, tandem duplication events. eLife 2020;9:e57947.

15. Khomarbaghi Z, Ngan WY, Ayan GB, Lim S, Dechow-Seligmann G, et al. Large-scale duplication events underpin population-level flexibility in tRNA gene copy number in Pseudomonas fluorescens SBW25. Nucleic Acids Res 2024;gkae049.

16. Silby MW, Cerdeño-Tárraga AM, Vernikos GS, Giddens SR, Jackson RW, et al. Genomic and genetic analyses of diversity and plant interactions of Pseudomonas fluorescens. Genome Biol 2009;10:R51.

17. Bertels F, Rainey PB. Within-genome evolution of REPINs: a new family of miniature mobile DNA in bacteria. PLoS Genet 2011;7:e1002132.

18. Bertels F, Rainey PB. Curiosities of REPINs and RAYTs. Mob Genet Elements 2011;1:262–268.

19. Reams AB, Roth JR. Mechanisms of Gene Duplication and Amplification. Cold Spring Harb Perspect Biol 2015;7:a016592.

20. Anderson P, Roth J. Spontaneous tandem genetic duplications in Salmonella typhimurium arise by unequal recombination between rRNA (rrn) cistrons. Proc Natl Acad Sci USA 1981;78:3113–3117.

21. Reams AB, Kofoid E, Savageau M, Roth JR. Duplication frequency in a population of Salmonella enterica rapidly approaches steady state with or without recombination. Genetics 2010;184:1077–1094.

22. Parab L, Romeyer Dherbey J, Rivera N, Schwarz M, Bertels F. Chloramphenicol reduces phage resistance evolution by suppressing bacterial cell surface mutants. 2023;2023.08.28.552763.

23. Deatherage DE, Barrick JE. Identification of mutations in laboratory-evolved microbes from next-generation sequencing data using breseq. Methods Mol Biol 2014;1151:165–188.

24. Pasternak C, Ton-Hoang B, Coste G, Bailone A, Chandler M, et al. Irradiation-induced Deinococcus radiodurans genome fragmentation triggers transposition of a single resident insertion sequence. PLoS Genet 2010;6:e1000799.

25. Shi Q, Parks AR, Potter BD, Safir IJ, Luo Y, et al. DNA damage differentially activates regional chromosomal loci for Tn7 transposition in Escherichia coli. Genetics 2008;179:1237–1250.

26. Nobusato A, Uchiyama I, Ohashi S, Kobayashi I. Insertion with long target duplication: a mechanism for gene mobility suggested from comparison of two related bacterial genomes. Gene 2000;259:99–108.

27. Weterings E, Chen DJ. The endless tale of non-homologous end-joining. Cell Res 2008;18:114–124.

28. Haviv-Chesner A, Kobayashi Y, Gabriel A, Kupiec M. Capture of linear fragments at a double-strand break in yeast. Nucleic Acids Research 2007;35:5192–5202.

29. Moore JK, Haber JE. Capture of retrotransposon DNA at the sites of chromosomal double-strand breaks. Nature 1996;383:644–646.

30. Ono R, Ishii M, Fujihara Y, Kitazawa M, Usami T, et al. Double strand break repair by capture of retrotransposon sequences and reverse-transcribed spliced mRNA sequences in mouse zygotes. Sci Rep 2015;5:12281.

31. Consuegra J, Gaffé J, Lenski RE, Hindré T, Barrick JE, et al. Insertion-sequence-mediated mutations both promote and constrain evolvability during a long-term experiment with bacteria. Nat Commun 2021;12:980.

32. Park SK, Mohr G, Yao J, Russell R, Lambowitz AM. Group II intron-like reverse transcriptases function in double-strand break repair. Cell 2022;185:3671-3688.e23.

33. King EO, Ward MK, Raney DE. Two simple media for the demonstration of pyocyanin and fluorescin. J Lab Clin Med 1954;44:301–307.

34. Romeyer Dherbey J, Parab L, Gallie J, Bertels F. Stepwise evolution of E. coli C and ΦX174 reveals unexpected lipopolysaccharide (LPS) diversity. Molecular Biology and Evolution 2023;40:msad154.

35. Ralser M, Querfurth R, Warnatz H-J, Lehrach H, Yaspo M-L, et al. An efficient and economic enhancer mix for PCR. BBRC 2006;347:747–751.

