## Supplementary Figure S1 for "Empirical evidence of a role for insertion sequences in the repair of DNA breaks in bacterial genomes"

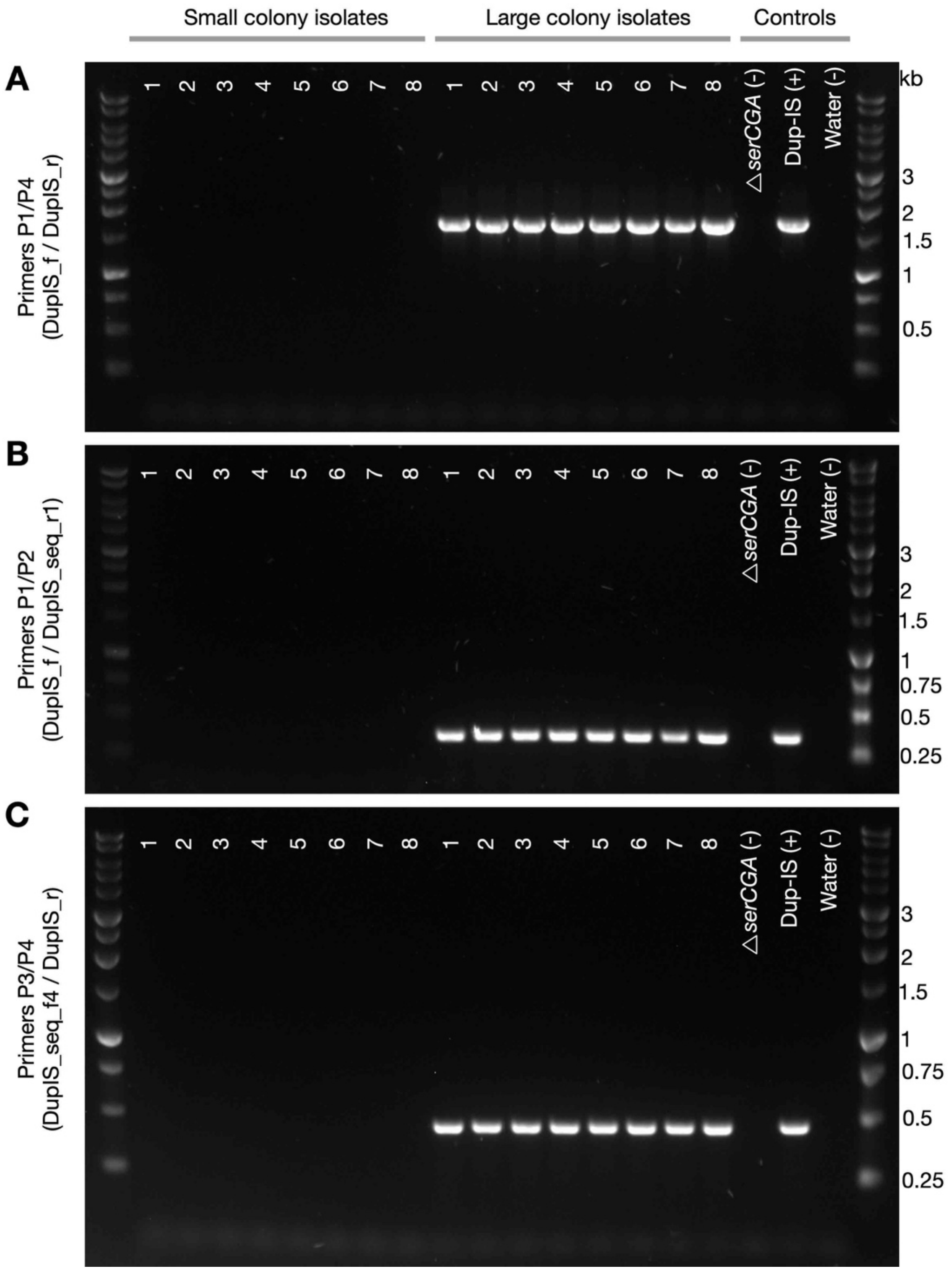


**Supplementary Figure S1. PCR results demonstrating the loss of the duplication fragment and the associated new IS*481* copy in all small colony isolates.** Agarose gel of small and large colony PCR products amplified with (**A**) external primers targeting both emergent junctions and the new IS*481* copy, (**B**) primers targeting junction Dup1-IS1, and (**C**) junction IS2-Dup2. Markers are 1-kb DNA ladders from Promega.
