## Supplementary Text S1 for "Empirical evidence of a role for insertion sequences in the repair of DNA breaks in bacterial genomes"

**Supplementary Text S1**
**The genome sequence of SBW25-Dup-IS**

### 1.1 The genomic rearrangement in SBW25-Dup-IS

Below is the predicted genomic sequence covering both emergent junctions and the new IS*481* copy in SBW25-Dup-IS. Note that the reported sequence is that of the leading strand (5'→3'), while the transposase is encoded on the lagging strand (5'→3').

>Dupcopy1_IS*481*_Dupcopy2 (leading strand, 5'→3')
…ACGCGCCGAAGCTGCCCTTCTCCTGCTCGATGTCGAGGATCATCTGCGCATTGCGCGGCACACTCTTGAGCTTG
CCCAGGTGGCGGATGATCCGCGTGTCCTGCATCAGCCGCTCCAGGTGCTCAGCGCCCATCAGCACGACCTGTAGA
ACCGTAGACATCGTTTACATCTGAAACCGGGGACATCGT**TTACACATTTGAGGCTTGGACGCGGGTCATACCTCG**
**TCCAAGCCTCCCCAAGGGATAATCACACAGGTACAAATCCCAGACATCATCCTCAGCTTCTTTGAGCCCTATCCT**
**TTCCCCGGATAATGCCTCGCTGACAAATACCAACTTGCCATTCCACTTGATCGATCCATCCTGCCTGACACTTCG**
**AACCCGCATTTCTGCCGGATACTCCACATCGGGTAAGCATCCTGGATAAAGTCGGGTAGACGGCATATACACATC**
**TCCCGGACGTTTCATACCAAGCGCTTCATGAGGGCGTACGTAATTGAACTCATGTTTGAAATGCTCCAGCAAAAG**
**CTGCTGCTCAACCAAATTGCTCCCTAGCGGTAGCTCAAGTTTCAGGCTGCGGTGCATTCGTTCGTGGCGACCATT**
**CTGGGCGGGCCTTCCTGGCATGGTTCGCTCGGGATAAATACCTAAACGGATCCACCAAACGGCCAGCGTGGACAT**
**TCTGGCCAGACCAGGAGAAGCGAAGGGGACGCCGTTGTCAGAGCGAATGACTTCCGGCATGCCGTACTCCTGAAA**
**AAGCCGCTCAAAGGTCTGCTTTACAGGTTGGGTCTTGATTTTAGGGTGAGCCCTGCAAGCCAGAATCAAACGGGA**
**TGCATGGTCTGTCACGGTCAGCGGGAAGCACATCTGCGCGTTGAGCATCTTGAATTGCCCTTTGTAGTCAGCGCA**
**CCAGGTTTTGTTGGGGTCATTGGCTTCTCGCATTTCAATATGAGAAGTGCCGTGCCGACGCTTGAATCGCCGCTT**
**ATTGACCAGCCCAAGCCGGTCAAGCCATTGGCCGGCTGTACTGGGAGACGGCCAATCGATAGAGGGATCTTCAAT**
**TCGCAATAGCTCGATGAGCTTCTTTGGCCCCCATTTATCGTGAGCCTCCTTCATGGCCACTACGCGAGCCAAGAT**
**CTCATCGTCGGTCTTGTTTGGGCTGTTATGAGGTCGCCGAGAAACCTCGGCTAACGACTTCAAATCGCCGTTATG**
**CCGAGAAATCCATTTGTCTACGGTCGGTCGACTGACACCGAAGCGGCGGGCCAGCTGGCTTTTCGTGAAATTACC**
**CGAAAGCCAGTCAGCAACCAGCTTGATTCGTTGGTTCATGGGGGACTCTTGGTTCCAGGGCAT**GATCAGTTACCT
CCTGATCATGCGTATTAACCTGTAAACCATGTCCCCGGTTAGAAATGTAAACGATGTCCCCGGTTTGTACAGGGC
AAGCCCCCTCCCACATTTGGATCGCGGCCTAACAGTTAGATGGTGGTCGGCTGTCAGGCAGCCGGCGGGAGCAAG
CTCCCTCGCCACAGGTCCCGCGCTATAGCCAGAATTGTGTAGATACCGATGCCCGATGGCGCTGAATCAGTCACC
TGATGAATCGACTGACACACCGCTATCGGGGGCAAGCCCCCTCCCACATTTGCATCGCGGTCTAACCGTTAGATG
GA…

- Blue: 3' region of duplication copy 1, located at 4,179,703-4,179,845 in SBW25 (non-repetitive, unique sequence within *pflu3785*)
- Yellow: the new 1279-bp IS*481* element; an exact match to three regions of SBW25 genome:
    - 2,338,017-2,339,292 (leading strand), *pflu2158* and surrounds
    - 4,792,869-4,794,144 (leading strand), *pflu4347* and surrounds
    - 6,387,109-6,388,384 (lagging strand), *pflu5832* and surrounds
- **Bold yellow: the 1149-bp transposase gene**
- Green: the 5' end of duplication copy 2, located at 4,119,905-4,120,133 in SBW25 (a highly repetitive intergenic region upstream of *pflu3723*).
- Green with turquoise highlighting: bases that render the sequence a unique match in the SBW25 genome.
- Underline: bases that match both sides of an emergent junction.

### 1.2 Computational evidence for the SBW25-Dup-IS genome rearrangement

SBW25-Dup-IS was sequenced using Illumina technology (150-bp paired-end reads). Raw reads were aligned to SBW25 reference sequence NC_012660.1 using *breseq* on standard settings [1]. The following summary statistics were produced:

| # raw reads | # mapped reads | % reads mapped | Genome coverage | |
|---|---|---|---|---|
| | | | Mean | St dev |
| 2,628,506 | 2,551,762 | 97.1 | 57.2 | 14.9 |

**Table 1.2.1**. Summary statistics of *breseq* alignment of the SBW25-Dup-IS Illumina whole genome re-sequencing data. Values are reported to 3 s.f.

Expectations for the SBW25-Dup-IS genome when aligned to the SBW25 reference:

i. Coverage of the duplication fragment (4,119,905 – 4,179,845) should be approximately two-fold that of the average genome coverage (due to the presence of 2 copies in SBW25-Dup-IS)
ii. Coverage of the IS*481* element should be approximately ~5-fold that of the average genome coverage (due to the presence of 5 copies in SBW25-Dup-IS)
iii. Emergent junctions Dup1-IS1 and IS2-Dup2 (see main manuscript Figures 1 and 3, Supplementary Figure S1) should be detectable.

**Points (i) and (ii): Coverage analyses**

Alignments of specific sequences with the raw reads were performed in Geneious Prime (version 11.1.4). The coverage analysis results are presented in Table 1.2.2 below. The results are consistent with SBW25-Dup-IS containing two copies of Dup and an extra copy of the IS*481* element.

| Genome coverage | | Duplication coverage[a] | | IS*481* coverage[b] | |
|---|---|---|---|---|---|
| Mean | St. dev | Mean | Normalized | Mean | Normalized |
| 57.2 | 14.9 | 113 | 1.98 | 282 | 4.93 |

**Table 1.2.2**. Sequencing data showing coverage across the duplicated region and the IS*481* element. [a]Coverage between reference sequence nucleotides 4,119,905 and 4,179,845 (the duplication fragment in SBW25-Dup-IS). [b]Coverage across the 1,279-bp IS*481* element (e.g., 4792868 - 4794146). Normalized coverage refers to the mean of interest divided by genomic mean. All values are reported to 3 s.f.

**Point (iii): Emergent junction analyses**

Below are two sequences (Dup1-IS1_30bp and IS2-Dup2_30bp), each crossing one of the emergent junctions in SBW25-Dup-IS. Both sides of each junction are covered by at least 15 bp. Neither junction sequence exists as a single stretch of 30 bp in the wildtype genome (or other non-SBW25-Dup-IS genomes), but both junction sequences are expected to exist in their entirety, once, within the derived SBW25-Dup-IS genome.

```
>Dup1-IS1_30bp
CCATCAGCACGACCTGTAGAACCGTAGACA

>IS2-Dup2_30bp
CCCCGGTTTGTACAGGGCAAGCCCCCTCCC
```

| # reads perfectly covering Dup1-IS1_30bp | # reads perfectly covering IS2-Dup2_30bp | Mean genome coverage (3 s.f.) |
|---|---|---|
| 51 | 55 | 57.2 |

**Table 1.2.3**. Data showing coverage across the two emergent junctions in the SBW25-Dup-IS genome. To be considered matching, a read must (i) cover all 30 bp of the junction in question, and (ii) match perfectly across the whole 30 bp (*i.e.*, no mismatches or gaps).

### 1.3 PCR based evidence for SBW25-Dup-IS genome rearrangement

In order to further test the bioinformatic predictions in the laboratory, we used the following PCR and Sanger sequencing protocol.

>Dup-IS_primers
…ACGCGCC**GAAGCTGCCCTTCTCCTGCTC**GATGTCGAGGATCATCTGCGCATTGCGCGGCACACTCTTGAGCTTG
CCCAGGTGGCGGATGATCCGCGTGTCCTGCATCAGCCGCTCCAGGTGCTCAGCGCCCATCAGCACGACCTGTAGA
ACCGTAGACATCGTTTACATCTGAAACCGGGGACATCGT**TTACACATTTGAGGCTTGGACGCGGGTCATACCTCG**
**TCCAAGCCTCCCCAAGGGATAATCACACAGGTACAAATCCCAGACATCATCCTCAGCTTCTTTGAGCCCTATCCT**
**TTCCCCGGATAATGCCTCGCTGACAAATACCAACTTGCCATTCCACTTGATCGATCCATCCTGCCTGACACTTCG**
**AACCCGCATTTCTGCCGGATACTCCACATCGGGTAAGCATCCTGGATAAAGTCGGGTAGACGGCATATACACATC**
**TCCCGGACGTTTCATACCAAGCGCTTCATGAGGGCGTACGTAATTGAACTCATGTTTGAAATGCTCCAGCAAAAG**
**CTGCTGCTCAACCAAATTGCTCCCTAGCGGTAGCTCAAGTTTCAGGCTGCGGTGCATTCGTTCGTGGCGACCATT**
**CTGGGCGGGCCTTCCTGGCATGGTTCGCTCGGGATAAATACCTAAACGGATCCACCAAACGGCCAGCGTGGACAT**
**TCTGGCCAGACCAGGAGAAGCGAAGGGGACGCCGTTGTCAGAGCGAATGACTTCCGGCATGCCGTACTCCTGAAA**
**AAGCCGCTCAAAGGTCTGCTTTACAGGTTGGGTCTTGATTTTAGGGTGAGCCCTGCAAGCCAGAATCAAACGGGA**
**TGCATGGTCTGTCACGGTCAGCGGGAAGCACATCTGCGCGTTGAGCATCTTGAATTGCCCTTTGTAGTCAGCGCA**
**CCAGGTTTTGTTGGGGTCATTGGCTTCTCGCATTTCAATATGAGAAGTGCCGTGCCGACGCTTGAATCGCCGCTT**
**ATTGACCAGCCCAAGCCGGTCAAGCCATTGGCCGGCTGTACTGGGAGACGGCCAATCGATAGAGGGATCTTCAAT**
**TCGCAATAGCTCGATGAGCTTCTTTGGCCCCCATTTATCGTGAGCCTCCTTCATGGCCACTACGCGAGCCAAGAT**
**CTCATCGTCGGTCTTGTTTGGGCTGTTATGAGGTCGCCGAGAAACCTCGGCTAACGACTTCAAATCGCCGTTATG**
**CCGAGAAATCCATTTGTCTACGGTCGGTCGACTGACACCGAAGCGGCGGGCCAGCTGGCTTTTCGTGAAATTACC**
**CGAAAGCCAGTCAGCAACCAGCTTGATTCGTTGGTTCATGGGGGACTCTTGGTTCCAGGGCAT**GATCAGTTACCT
CCTGATCATGCGTATTAACCTGTAAACCATGTCCCCGGTTAGAAATGTAAACGATGTCCCCGGTTTGTACAGGGC
AAGCCCCCTCCCACATTTGGATCGCGGCCTAACAGTTAGATGGTGGTCGGCTGTCAGGCAGCCGGCGGGAGCAAG
CTCCCTCGCCACAGGTCCCGCGCTATAGCCAGAATTGTGTAGATACCGATGCCCGATGGCGCTGAATCAGTCACC
TGATGAATCGACTGACACACCGCTATCGGGGGCAAGCCCCCTCCCACATTTGCATCGCGGTCTAACCGTTAGATG
GAGCGGGAGCAAACGCCTACACGTCACAACAGTGTCCGAGCCAGCGTCTTTTTCCACGCTGAATAACGCCCCGCC
ATCGCG**CAATCATCCTGCGGCTCAAACCG**CTCCCGCTGCC…

Primer set 1 (main manuscript Figure 1D): targeting both emergent junctions and the new IS copy between the two junctions.

DupIS_f **GAAGCTGCCCTTCTCCTGCTC**
DupIS_r **CGGTTTGAGCCGCAGGATGATTG**

Expected product sizes:
SBW25-Dup-IS: 1,746 bp
Other strains (e.g., SBW25, SBW25Δ*serCGA*): no product

Additional primers to sequence full-length SBW25-Dup-IS PCR product (a amplify junctions):
DupIS_seq_r1: **CAGGCAGGATGGATCGATCAAG**
DupIS_seq_f1: **CTTGATCGATCCATCCTGCCTG**
DupIS_seq_f2: **GTTGTCAGAGCGAATGACTTCC**
DupIS_seq_f3: **GTACTGGGAGACGGCCAATC**
DupIS_seq_f4: **CAACCAGCTTGATTCGTTGGTTC**
