## Supplementary Text S2 for "Empirical evidence of a role for insertion sequences in the repair of DNA breaks in bacterial genomes"

### Supplementary Text S2
### The genome sequence of Ec-Del-IS

#### 2.1 The genomic rearrangement in Ec-Del-IS

- A 62,007-bp deletion (3,612,369 – 3,674,375 inclusive), plus a new IS*5*/IS*1182* element copy in place of the deleted fragment.
- There are 3 identical copies of IS element in Ec-Del-IS, and two identical (complete) copies in the wildtype *E. coli* C genome.

>Ec-Del-IS<br>
TGCCGGACCGCCGACC**ACATACACGCCACAGGAGG**CGTTATCCGCCACGGTATCAACAAATTTGATCGACCAGTC<br>
GCGGATACGGCTGCCGACCACTTCCAGCGCGGGAAGTACCCACTCTATGGCGTTGTAAAGCTCGTCAAAAGGGCG<br>
TTGTTGCTCAAATCAGGGAACTATCCCGCAGTTTGGTGATCTGATCTCTTAGGTTCACCACCGGGAGCGCCAGAT<br>
GGGCAAGTCCAGGCACAAGATCAGCAACTGGAAACAGTACAACCAAGCCTTGGTGCAACGTGGTTCGTTGACCGT<br>
CTGGATGGATGAGCAAGCCATCCAGCAGTGGCATTGCCAAACCCATCATGGCCGCCGAGGG**CGAGGCTTCCATTA**<br>
**CAGC**GACTCCGCCATTGAAACCGCGCTGATGCTCAAGGCGGTATTCAAGCTACCGCTGCGTGCCCTGGAAGGCTT<br>
CATCAACTCGCTGTTCAAGCTGATGAAGGTGCCCTTACAGTCGCCGGACTACAGCTGTATCAGCAAGCGCGCCAA<br>
GACGGTCGAAATCAAGTACCGCCTACCGAGCCATGGCCCAGTGGCTCACCTGGTCATCGATGCCACCGGACTCAA<br>
GGTCTATGGCGAAGGCGAATGGAAAATTCGCAAGCACGGCAAGGAGAAACGCCGCGTCTGGCGCAAGCTTCACCT<br>
GGCCGTAGATGCGG**CGACCCATGCCATCGTTG**CAGCCGAAGTCAGCCTGGAAACGGTGGGCGATAACGAGGTGTT<br>
GCCCACGCTGCTCAACCCCTTGCGCCGCAAGATAGAACAAGTCAGCGCCGATGGTGCCTATGACACCAGAGCCTG<br>
TTATGCCCTACTGCAAAAGAAGGGCGCCAAGGCCACCATACCGCCGAGAAAAAATGCGGCATTCTGGAAGACAGG<br>
CCATCCACGCAACGAGGCGGTTGCCGCGCTCAAGGCAGGAGAACTGGAGCAATGGAAAAAGG**ACTCTGGCTATCA**<br>
**TCAGCGC**TCGATAGCTGAGACCGCCATGTATCGCTTCAAGCAGCTCATCGGTCCGAAACTGAGTCTGCGGAGCTA<br>
CAACGCCCAGGTGGGTGAGATCCTGGCCGGGGTGAAAGTCATGAACAAGGTCATAGGGCTTGGTATGCCTATTCG<br>
GCAGGCTGTCAACTAAGCCCGCCCGGCGGGTTGGG**GAGCGACCATCCAACTTTG**GATTTGATCAACAAGGCCTTT<br>
AGACACCCAAAAAGAGAACCGCCCCAAATGTGTGACATATCTGGATCTATTTGTGCAGTTTTACATAACATCATA<br>
CACCGCTCATGTATTTCTCCGCAGTCACTATCGTATGCTGTTTGGCAGGTTAATAAATATCCAGATGATTCCCCG<br>
TTACAAGACGAAATTGTCGATTCCATCTTACTTTTAAGTAATGCGGATAAGTTTTGCATTGACAATGCTGGAGCC<br>
ATAACAGATAGCCCCAT**CGGATCAATGTTCATAACCGGACTC**AACGGATAAATGTATAAATTCCATCCCCCCTCC<br>
A

BLUE end of gene 1 (*B6N50_18465*, encoding a 2-keto-4-pentenoate hydratase); this is the region where the deletion fragment starts (i.e., Junction 1: Del1-IS1)<br>
BLACK IS element sequence (1,051-bp)<br>
*ITALICS* transposase gene (918-bp)<br>
GREEN single nucleotide from gene 2 (*B6N50_18845*, encoding a hypothetical protein) at Junction 2 (IS2-Del2)<br>
GREEN intergenic region at end of deletion (i.e., Junction 2: IS2-Del2)<br>
PURPLE gene 3 (*B6N50_18850*, encoding a hypothetical protein), after the intergenic region downstream of junction 2 (IS2-Del2)

30-bp Junction 1 sequence: "Del1-IS1": AAAGCTCGTCAAAAGGGCGTTGTTGCTCAA (grey = side 1, turquoise = side 2)<br>
30-bp Junction 2 sequence: "IS2-Del2": TTGATCAACAAGGCCTTTAGACACCCAAAA (grey = side 1, green / turquoise = side 2)

#### 2.2 PCR based evidence for Ec-Del-IS genome rearrangement

##### *2.2.1 PCR to amplify the entire rearrangement region*

DelIS_LP_F: **ACATACACGCCACAGGAGG**<br>
DelIS_LP_R: **GAGTCCGGTTATGAACATTGATCCG**

Expected product sizes:<br>
wild type: ~62 kb (i.e., too large to amplify here)<br>
Ec-Del-IS: 1,451 bp

Deletion without new IS element copy: ~400 bp (as far as we know, this doesn't exist)

Additional primers to sequence through PCR product:
`DelIS_LP_seq_F1:` **CGAGGCTTCCATTACAGC**
`DelIS_LP_seq_R1:` **GCTGTAATGGAAGCCTCG** `(reverse complement of DelIS_LP_seq_F1)`
`DelIS_LP_seq_F2:` **CGACCCATGCCATCGTTG**
`DelIS_LP_seq_F3:` **ACTCTGGCTATCATCAGCGC**
`DelIS_LP_seq_F4:` **GAGCGACCATCCAACTTTG**

##### *2.2.2 PCRs for amplifying each emergent junction*

**Junction 1 (Del1-IS1)**
`DelIS_LP_F:` **ACATACACGCCACAGGAGG**
`DelIS_LP_seq_R1:` **GCTGTAATGGAAGCCTCG**

Expected product sizes:
wild type: no product expected
Ec-Del-IS: 363 bp
Deletion without new IS element copy: no product expected

**Junction 2 (IS2-Del2)**
`DelIS_LP_seq_F4:` **GAGCGACCATCCAACTTTG**
`DelIS_LP_R:` **GAGTCCGGTTATGAACATTGATCCG**

Expected product sizes:
wild type: no product expected
Ec-Del-IS: 307 bp
Deletion without new IS element copy: no product expected

##### *2.2.3 Probe PCR for amplifying a wild-type-specific product*

`DelIS_LP_F:` **ACATACACGCCACAGGAGG**
`DelIS_WT_LP_R` **GAAGCCGAAATTGCGCTGGTG** `(binds in WT genome but not Ec-Del-IS)`

Expected product sizes:
wild type: 182 bp
Ec-Del-IS: no product expected
Deletion without new IS element copy: no product expected

```
>Ecoli_C_WT_genome
…ACATACACGCCACAGGAGGCGTTATCCGCCACGGTATCAACAAATTTGATCGACCAGTCGCGGATACGGCTGCC
GACCACTTCCAGCGCGGGAAGTACCCACTCTATGGCGTTGTAAAGCTCGTCAAAAGTGGTGTCGGTATTGGGCAG
ATCGCGGTTCAGCACCAGCGCAATTTCGGCTTCAATACGCGGTTGCAGCACACGGGAAAACGGATGGTTTCGTT
ATCGCCATAACACATATCAGCAAACAGCGTACCGAAGTCCGGTTGATCGACACCCAGCTGTTGCTGCACTTTCGG
GTGCGTCAG…
```

`Yellow` deleted in Ec-Del-IS
